# Mice generated with induced pluripotent stem cells derived from mucosal-associated invariant T cells

**DOI:** 10.1101/2022.07.27.501791

**Authors:** Chie Sugimoto, Hiroyoshi Fujita, Hiroshi Wakao

## Abstract

The function of mucosal-associated invariant T (MAIT) cells, a burgeoning member of innate-like T cells abundant in humans and implicated in many diseases, remains obscure. To explore this, mice with a rearranged T cell receptor (TCR) α or β locus, specific for MAIT cells, were generated via induced pluripotent stem cells derived from MAIT cells and designated Vα19 and Vβ8 mice, respectively. Both mice expressed large amounts of MAIT cells. The MAIT cells from these mice were activated by cytokines and an agonist to produce IFN-γ and IL-17. While Vβ8 mice showed resistance in a cancer metastasis model, Vα19 mice did not. Adoptive transfer of MAIT cells from the latter into the control mice, however, recapitulated the resistance. This has implications for understanding the role of MAIT cells in health and disease and in developing treatments for the plethora of diseases in which MAIT cells are implicated.

## Introduction

MAIT cells belong to an emerging member of innate-like T cells. They are characterized by the semi-invariant T cell antigen receptor (TCR), dependence on major histocompatibility complex (MHC) I-related gene protein (MR1), on microbiota for their development, expression of the transcription factors promyelocytic leukemia zinc finger protein (PLZF), and retinoic acid receptor-related orphan receptor gamma t (RORγt) (Franciszkiewicz et al., 2016, Godfrey et al., 2019, Treiner et al., 2003). PLZF is associated with tissue homing and residency, conferring a rapid antigen response to T cells in a TCR-independent manner, thus essential for development and function of MAIT cells (Beaulieu and Sant’Angelo, 2011, Koay, Hui-Fern et al., 2016). RORγt is responsible for T helper (Th)-17 cell differentiation and for the development of type 3 innate lymphoid cells (ILC3) (Eberl, 2017). Moreover, MAIT cells recognize small compounds, such as vitamin B2 metabolites, B9, and derivatives thereof, presented on MR1 via the semi-invariant T cell receptor (TCR) α (TRAV1-2-TRAJ33 (-TRAJ12, -TRAJ20)) for human and TRAV1-TRAJ33 for mouse, paired with a limited set of TCR β repertoire, as is the case for other innate-like T cells (Tilloy et al., 1999, Kjer-Nielsen et al., 2012, Corbett et al., 2014). Whilst T cells, in adaptive immunity, necessitate priming and clonal expansion for exerting their effector functions, innate-like T cells do not. This is primarily due to the abundance of the latter, which are poised to respond to stimuli *in vivo*. MAIT cells are implicated in an array of diseases, such as bacterial and viral disease, autoimmune, inflammatory, and metabolic diseases, asthma, and cancer (Godfrey et al., 2019, Godfrey et al., 2015, Toubal et al., 2019). However, the precise role of MAIT cells in each disease is enigmatic despite the abundance of MAIT cells in humans, representing 1-10% of T cells of the peripheral blood mononuclear cells (PBMC) and 20-45% in the liver (Dusseaux et al., 2011).

Our recent study established induced pluripotent stem cells (iPSCs) from murine MAIT cells (MAIT-iPSCs). MAIT-iPSCs exclusively gave rise to MAIT-like cells as defined 5-(2-oxopropylideneamino)-6-D-ribitylaminouracil (5-OP-RU)-loaded mouse MR1-tetramer (mMR1-tet)^+^TCRβ^+^ cells, when differentiated under T cell permissive culture conditions. These MAIT-like cells shed light on the possible roles of MAIT cells in cancer immunology (Sugimoto et al., 2022). Previously, we established cloned mice and cloned embryonic stem (ntES) cells by nuclear transfer of iNKT cells, which are another member of the innate-like T cells (Inoue et al., 2005, Wakao et al., 2008). The cloned mouse offspring with rearranged TCRα specific for iΝΚΤ cells (*Trav11-Traj18*) showed an enhanced frequency of iNKT cells (Wakao et al., 2007), and the ntES cells exclusively redifferentiated into iNKT-like cells in the OP9 and OP9/(delta-like 1) DLL1 system (Wakao et al., 2008). Therefore, we reasoned that mice with rearranged TCRα and/or β specific for MAIT cells would have an increased frequency of MAIT cells. Given that iPSCs possess pluripotency, including the ability to give rise to chimeric mice as ES cells, we explored the ability of MAIT-iPSCs to confer the rearranged TCR *loci* to the mice (Takahashi and Yamanaka, 2006).

In this study, we generated mice from MAIT-iPSCs with the rearranged TCRα or β locus, specific for MAIT cell TCR, designated as Vα19 and Vβ8 mice, respectively. TCR repertoire analysis on *Trav* in Vβ8 mice revealed more diversity in the CDR3α domain than anticipated. While both mice possessed increased MAIT cell numbers, Vβ8 mice exhibited resistance to the inoculated cancer, but Vα19 mice did not. Intriguingly, adoptive transfer of MAIT cells from Vα19 mice into the control mice conferred a similar resistance. Further study identified T cells, in particular, CD8 T cells as an adjuvant for MAIT cells to exert anti-tumor immune functions.

## Results

### Mice harboring rearranged TCR loci specific for MAIT cells

We previously established several iPSC clones derived from 5-OP-RU-loaded mouse MR1 tetramer (mMR1-tet) positive MAIT cells and confirmed that some clones possessed pluripotency through chimera formation (Sugimoto et al., 2022). Using the chimeric mice, we then attempted to generate novel C57BL/6 (B6) strains harboring rearranged TCR loci specific for MAIT cells. Three MAIT-iPSC clones, L7-1, L11-1, and L19-1 were injected independently into ICR 8-cell stage embryos, resulting in 11 chimeric male mice with varying degrees of chimerism (<30-90% as defined by donor coat color) (Table 1). Mice with 60∼90% chimerism were crossed with C57BL/6 females and resulting black-colored pups were screened for the presence of the rearranged configuration for *Trav1-Traj33* and/or *Trbv* corresponding to the original iPSC clones (Figure S1). Among the chimeric mice, one line with 60-90% chimerism (#26 derived from iPSC clone L7-1 harboring *Trav1-Traj33* and *Trbv8-2-D1-J1-2* (Sugimoto et al., 2022)) successively gave rise to offspring bearing *Trav1-Traj33, Trbv8-2-D1-J1-2*, or both (Table 1, Figure S1). Hereafter, these mouse strains are labeled Vα19 (harboring *Trav1-Traj33*), Vβ8 (harboring *Trbv13-3-d1-j1-2*), and Vα19/Vβ8 (harboring both *Trav1-Traj33* and *Trbv13-2-d1-j1-2*), respectively.

**Table 1.**
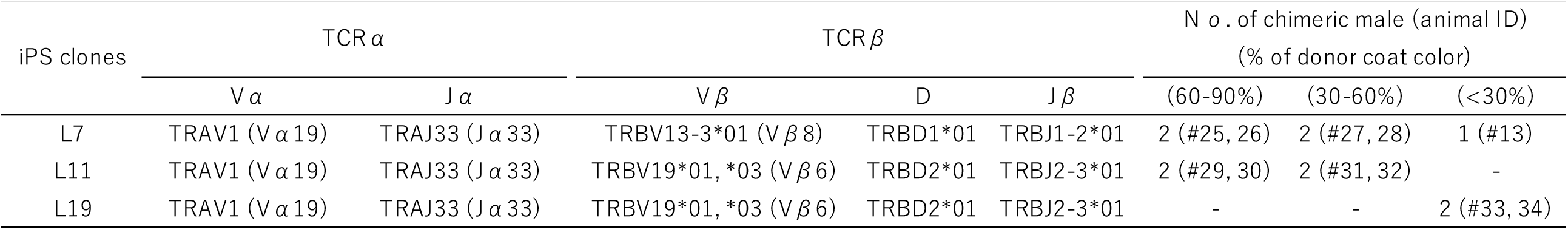
Characterization of iPSCs and chimeric mice used for germline transmission. The iPSC clones used for chimeric mouse generation are indicated in the left column with the corresponding usage of Vα, Jα, Vβ, Dβ, and Jβ. The number of chimeric mice harboring 60-90% of the chimerism is shown with the proper identifier (#) in the right column.

To visualize the impact of the rearranged TCRα or TCRβ locus specific for MAIT cells on the frequency of MAIT cells, flow cytometric analysis was performed using peripheral blood (Figure 1). While the frequency of MAIT cells (defined as mMR1-tet^+^TCRβ^+^ cells) in the control mice rarely exceeded 0.1% among αβ T cells, in Vβ8 mice and Vα19 mice it reached ∼0.5% and ∼36%, respectively. Furthermore, the presence of *Trav1-Traj33* and *Trbv8-2-D1-J1-2* enhanced the frequency (Figure 1A and B). CD4 and CD8 expression analysis identified an increase in CD4^-^CD8^-^ MAIT cells concomitant with a decrease in CD4^+^ MAIT cells, relative to the control mice (Figure 1C). In contrast, among mMR1-tet^-^TCRβ^+^ cells (non-MAIT T cells) in Vβ8 mice, there was little change in the frequency of CD4^+^ and CD8^+^ as well as CD4^-^CD8^-^ T cells. However, relative to the control, the frequency of CD4^-^CD8^-^ T cells in Vα19 mice increased, and, in Vα19/ Vβ8 mice, CD8^+^ T cell frequency decreased (Figure 1D). Hereafter, all the experiments were performed with Vα19 and Vβ8 mice.

**Figure 1.**
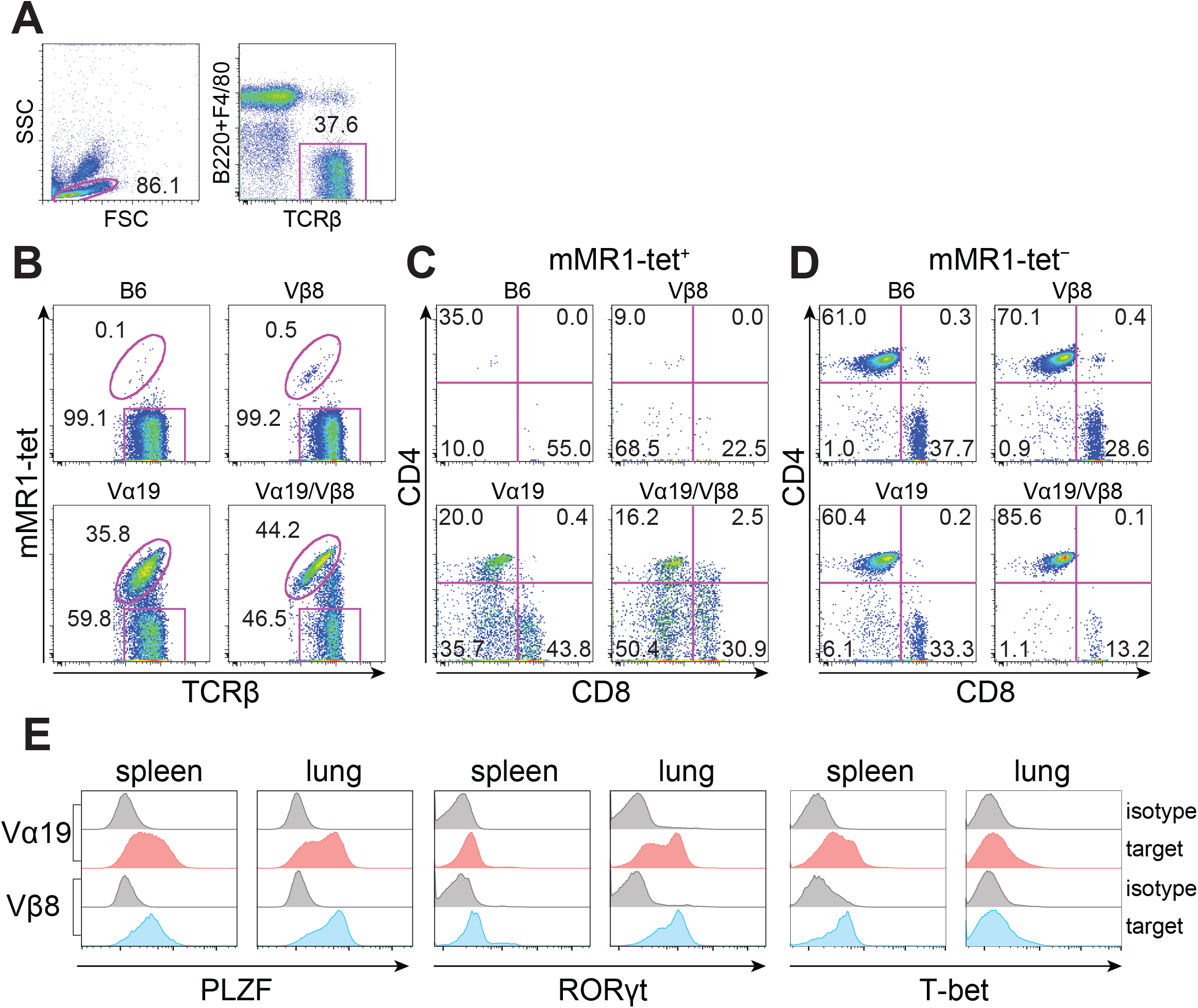
Blood MAIT cell profile in C57BL/6 (control), Vα19, Vβ8, and Vα19/ Vβ8 mice. Anticoagulated mouse peripheral blood collected by submandibular bleeding was used for flow cytometric analysis. (A) Gating for T cell population (TCRβ^+^ lymphocytes). (B) Analysis of MAIT cells (mMR1-tet^+^TCRβ^+^) and non-MAIT T cells (mMR1-tet^-^TCRβ^+^) in each mouse strain. (C and D) The expression of CD4 and CD8 in mMR1-tet^+^ TCRβ^+^ cells (C) and mMR1-tet^-^TCRβ^+^ cells (D). B6: C57BL/6, Vβ8: Vβ8 mice, Vα19: Vα19 mice, Vα19/ Vβ8: Vα19/ Vβ8 mice. The number shows the percentage of the indicated cell subset. Representative data from three independent experiments are shown. (E) Representative histograms showing the expression of PLZF, RORγt, and T-bet in Vα19 and Vβ8 mouse spleen and lung MAIT cells (shaded in pink and blue, respetively). Isotype control staining is also indicated (shaded in grey). Representative data from three independent experiments are shown.

PLZF, RORγt and T-bet are transcription factors which play a pivotal role in development and function of MAIT cells (Koay et al., 2016, Cui et al., 2015, Rahimpour et al., 2015). We then examined those expression in MAIT cells from the spleen and lungs of Vα19 and Vβ8 mice. PLZF was detected in majority of spleen and lung MAIT cells in both Vα19 and Vβ8 mouse (Figure 1 E). RORγt is also detected in MAIT cells from both mice. However, lung MAIT cells showed higher expresson than spleen MAIT cells (Figure 1 E).

MAIT cells are composed of MAIT1 and MAIT17 subset, however, their distribution varies from one tissue to another in C57BL/6 (Legoux et al., 2020). MAIT17 is characterized with RORγt expression, whereas MAIT1 is characterized with T-bet expression. We then evaluated the relative abundance of MAIT1 and MAIT17 in these mice. In both Vα19 and Vβ8 mouse lung MAIT cells, RORγt-expressing cells (MAIT17) were dominant over T-bet-expressing cells (MAIT1) (∼60-80% cells were RORγt^+^, while less than 20% cells were T-bet^+^). In contrast, MAIT1 was more abundant in the spleen MAIT cells (40 - 60 % of cells were T-bet^+^, while less than 10% and 40% were RORγt^+^ in Vα19 mice and Vβ8 mice, respectively (Figure S2). The results were in line with the previous report with C57BL/6 (Legoux et al., 2020).

Given the elevated frequency of MAIT cells in the blood, we next sought whether Vα19 and Vβ8 mice also harbored an enhanced frequency of MAIT cells across the organs. In Vα19 mice, MAIT cells constituted ∼20-25% of CD3^+^ T cells in the spleen, inguinal and mesenteric lymph nodes, intestinal lamina propria (LP), and the thymocytes, while increased MAIT cell numbers were observed in the liver, lung, and bone marrow (∼35-40% among CD3^+^ T cells) (Figure 2). In Vβ8 mice, albeit to a lesser extent than Vα19 mice, MAIT cells represented 2-10% of CD3^+^ T cells in the analyzed tissues (Figure 2A). Finally, the control mice exhibited a much lower MAIT cell frequency.

**Figure 2.**
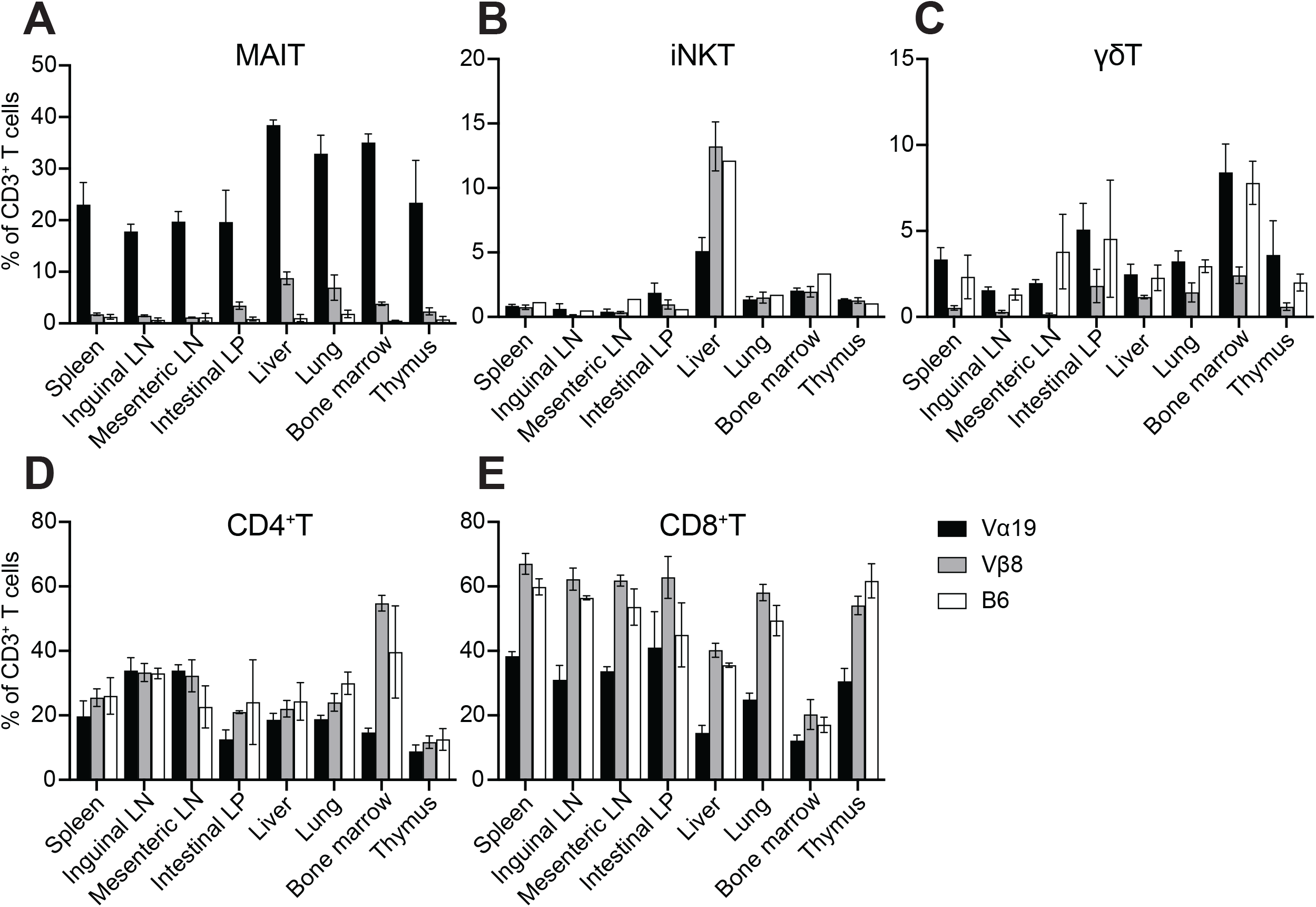
Frequency of MAIT, iNKT, γδT, CD4^+^ T, and CD8^+^ T cells in Vα19, Vβ8, and control mice. Relative frequency of each subset among CD3^+^ T cells in the indicated organs is shown (n=4 per strain). Bar graph depicts mean±SD. Vα19: Vα19 mice, Vβ8: Vβ8 mice, B6: C57BL/6

As *Trav1* is located at the most upstream of the *Trav* locus, rearranged *Trav1*-*Traj33* is unlikely to undergo further rearrangement (secondary rearrangement) in the same allele. We thus evaluated whether *Trav1*-*Traj33* impacted the generation of other T cells such as innate-like T cells and T cells in adaptive immunity (Figures 2 and S3). The frequency of iNKT cells, the most abundant mouse innate-like T cell subset, was reduced in the liver of Vα19 mice relative to the control mice, while no difference was apparent in Vβ8 mice (Figure 2B). In sharp contrast, the frequency of γδT cells decreased in Vβ8 mice, but Vα19 showed little difference compared with the control mice (Figure 2C). While the frequency of CD4^+^ T cells among Vα19, Vβ8, and control mice did not differ across the tissues, except for a decrease in the bone marrow of Vα19 mice, that of CD8^+^ T cells in Vα19 mice tended to decline in all tissues examined relative to other mouse strains (Figure 2D and E).

Analysis of myeloid and lymphoid cells identified a decrease in T cell frequency in the intestinal LP and lung lymphocytes among CD45^+^ cells in Vα19 mice relative to the other strains. Nonetheless, the frequency of B cells (CD45^+^CD11b^-^CD3^-^CD19^+^ cells), NK cells (CD45^+^CD11b^-^CD3^-^CD19^-^NK1.1^+^NKG2D^+^ cells), macrophages (CD45^+^ CD11b^dim^), neutrophils (CD45^+^CD11b^+^Ly6G^+^ cells), and monocytes (CD45^+^CD11b^+^Ly6C^+^ cells) was unaffected in both Vα19 and Vβ8 mice (Figure S4).

In summary, the data demonstrated that the rearranged TCR configuration specific for MAIT cells affected T cell development, and resulted in an enhanced frequency of MAIT cells in mice.

### Expression of the molecules relevant to MAIT cells

Comparative analysis of MAIT cells and of non-MAIT T cells in the representative tissues of Vα19 and Vβ8 mice was performed based on the cell surface molecules (Cui et al., 2015, Rahimpour et al., 2015). In Vα19 mice, little difference in surface molecule expression was revealed between MAIT and non-MAIT T cells of any molecules except for NK1.1 in the spleen and in the intestinal LP, while there was an increase in interleukin (IL)-18Ra and CXCR6 in the liver and in lung MAIT cells relative to non-MAIT T cells (Figure 3). In contrast, between MAIT cells and non-MAIT T cells of Vβ8 mice, there was a large difference in surface-molecule expression levels (Figure 3). MAIT cells in Vβ8 mice expressed higher levels except for CD62L, which was expressed less than non-MAIT T cells across the tissues, except for intestinal LP. CD103 showed a similar decrease in MAIT cells in intestinal LP, whilst the reverse trend was seen in the spleen, liver, and lung in Vβ8 mice (Figure 3).

**Figure 3.**
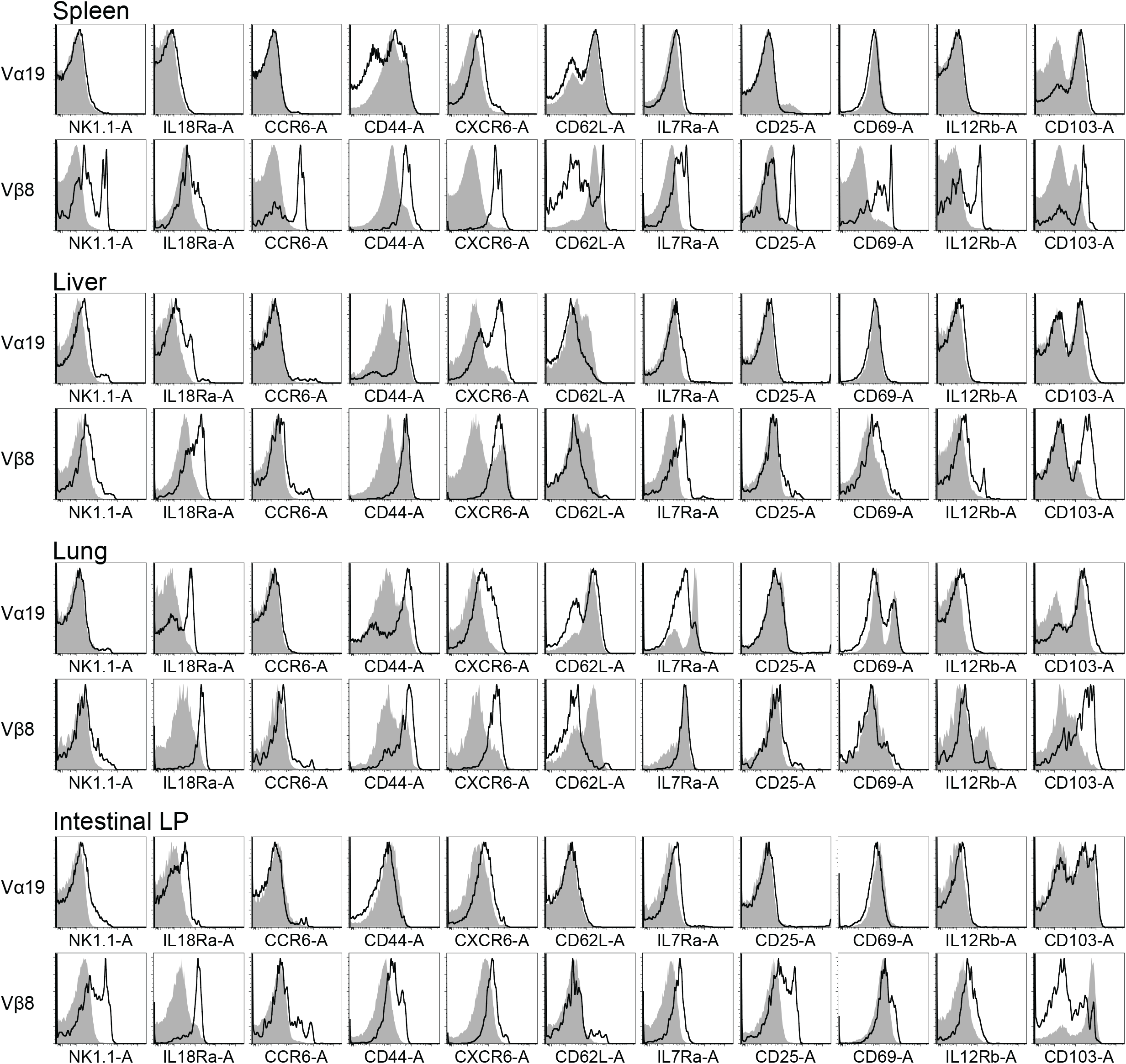
Comparative phenotypic analysis of MAIT cells and non-MAIT T cells. Expression levels of the indicated markers in MAIT cells (thick black lines) and in non-MAIT T cells (shaded grey) from the spleen, liver, lung, and intestine of Vα19 and Vβ8 mice are depicted as a histogram. Data are representative of three independent experiments.

### TCR repertoire in Vα19 and Vβ8 mouse

Though the rearranged *Trav19-Traj33* and/or *Trvb13-3-d-j* enhanced the frequency of MAIT cells, the impact of such gene rearrangement on TCR repertoire in the thymus was unknown. We thus analyzed the TCR repertoire in Vα19 and Vβ8 mouse thymi. MAIT cells (mMR1-tet^+^TCRβ^+^ cells) and non-MAIT T cells (mMR1-tet^-^CD4^+^CD8^+^TCRβ^+^) were sorted from the thymocytes and subject to high-throughput sequencing. While TCRα chains in MAIT cells were mostly composed of *Trav1-Traj3*3 in both mouse strains (94.6% in Vα19 mice and 85.1% in Vβ8 mice), other combinations such as *Trav16D-Traj18* (2.2%), *Trav1-Traj30* (0.3%), *-31*(0.3%), *-27* (0.1%), besides *Trav1-Traj12* (0.1%) were also found in Vβ8 mouse (Figure 4A and Table S1) (Reantragoon et al., 2013, Koay, H-F et al., 2019). Further analysis of *Trav1-Traj3*3 revealed that TCRα complementarity determining region 3 (CDR3α) sequences were quasi-invariable in Vα19 mice, whereas in Vβ8 mice they were highly divergent both in length and in sequence (Figure 4C) (Rahimpour et al., 2015).

**Figure 4.**
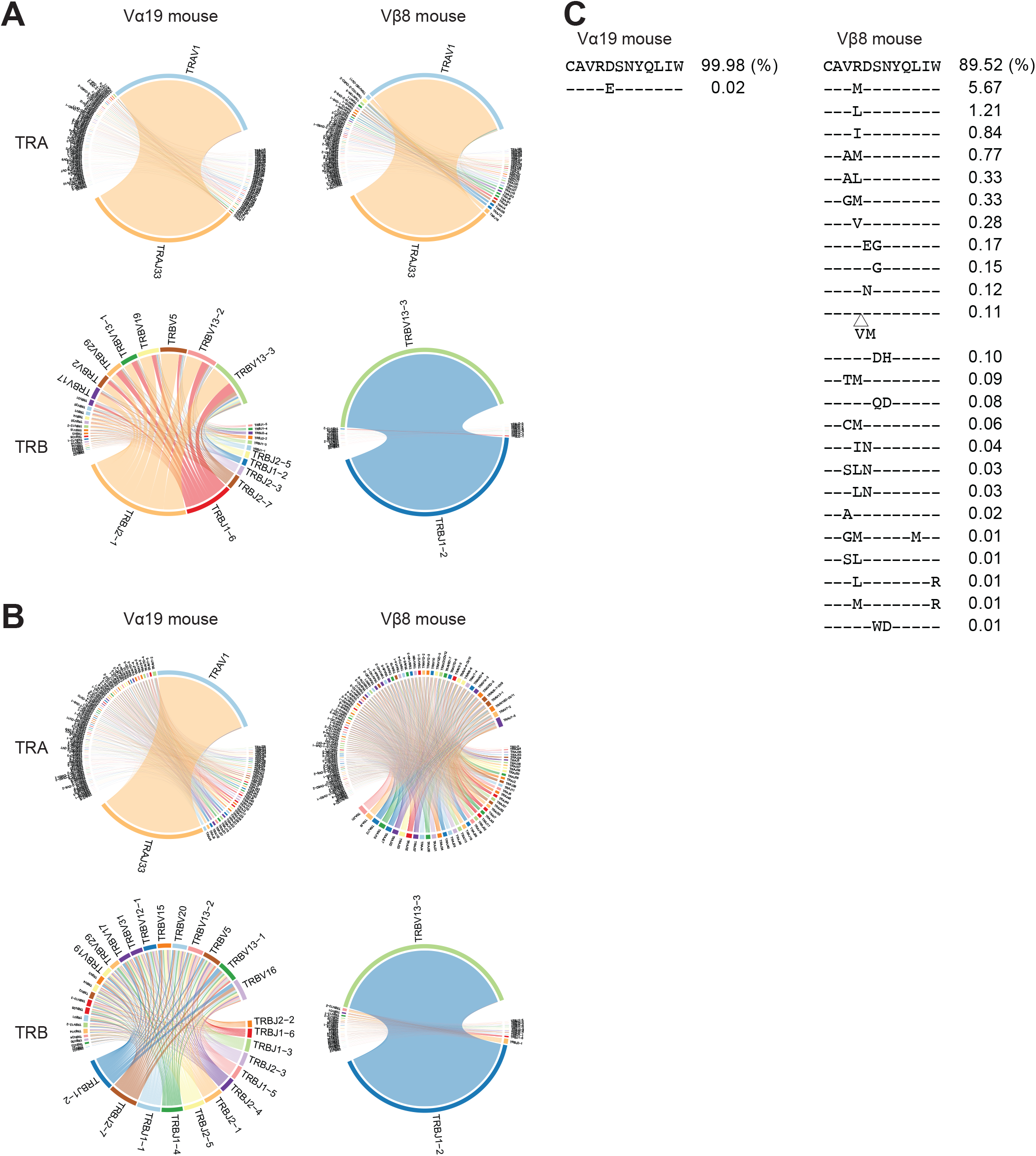
TCR repertoire analysis. (A) Circos plot for MAIT cells. Circos plots for MAIT cells in the thymocytes from the indicated mouse strain are depicted. Upper and lower panels indicate TRA and TRB, respectively. (B) Circos plot for non-MAIT T cells. Circos plots for non-MAIT T cells in the thymocytes from the indicated mouse strain are shown. Upper and lower panels indicate TRA and TRB, respectively. (C) Diversity in CDR3α. Amino acid sequences of CDR3α are aligned for MAIT cells in Vα19 mouse (left panel) and those in Vβ8 mice (right panel). -; the same amino acid indicated in the first line of sequences. .△; additional sequence insertion point. The percentage shows the frequency of the clone(s) harboring the indicated CDR3 sequence among MAIT cells.

TRBV repertoire usage was quasi-exclusively limited to *Trbv13-3-Trbj1-2* in Vβ8 mouse MAIT cells. In contrast, no such exclusive usage was seen in Vα19 mice TRBV repertoire, although there existed biased use of *Trbv13-3, 13-2, 5, 19*, and *13-1* (Figure 4A). As expected, CDR3β sequences of *Trbv13-3-Trbj1-2* in Vβ8 mouse MAIT cells exclusively comprised a single clone, while those for Vα19 mouse MAIT cells were divergent (Table S3).

TRAV repertoire of non-MAIT T cells in Vα19 mice showed enrichment in *Trav1-Traj3*3 similar to MAIT cells, while that in Vβ8 mice represented much more diversity. Regardless of the enrichment in *Trav1-Traj3*3 in non-MAIT T cells, TRBV repertoire was less biased in Vα19 mice. In sharp contrast, *Trbv13-3-Trbj1-2* was always predominant in Vβ8 mice, even in non-MAIT T cells (Figure 4B) (Rahimpour et al., 2015).

### MAIT cell activation and production of IFN-γ and IL-17A

MAIT cells are activated to produce cytokines such as IFN-γ and IL-17A in a TCR dependent and/or independent manner (Jesteadt et al., 2018, Leng et al., 2019, Rahimpour et al., 2015, Hinks and Zhang, 2020). Our recent study also found that redifferentiated MAIT cells from MAIT-iPSCs *in vitro* possessed similar properties (Sugimoto et al., 2022). Therefore, cells isolated from the spleen and the lung of Vα19 and Vβ8 mice were first stimulated with varying concentrations of the MAIT cell agonist 5-OP-RU, and the resulting activation through TCR was evaluated by CD69 and CD25 expression (Figure 5A and Figure S5). Similarly, while activation of lung and spleen MAIT cells from Vα19 mice was observed in a 5-OP-RU dose-dependent manner, that from Vβ8 mice reached plateau at 10 nM, resulting in less enhanced activation (Figure 5A). Notably, Vβ8 mouse lung MAIT cells already showed a degree of CD25 and CD69 expression in the absence of stimulation (Figure 5A and Figure S5).

**Figure 5.**
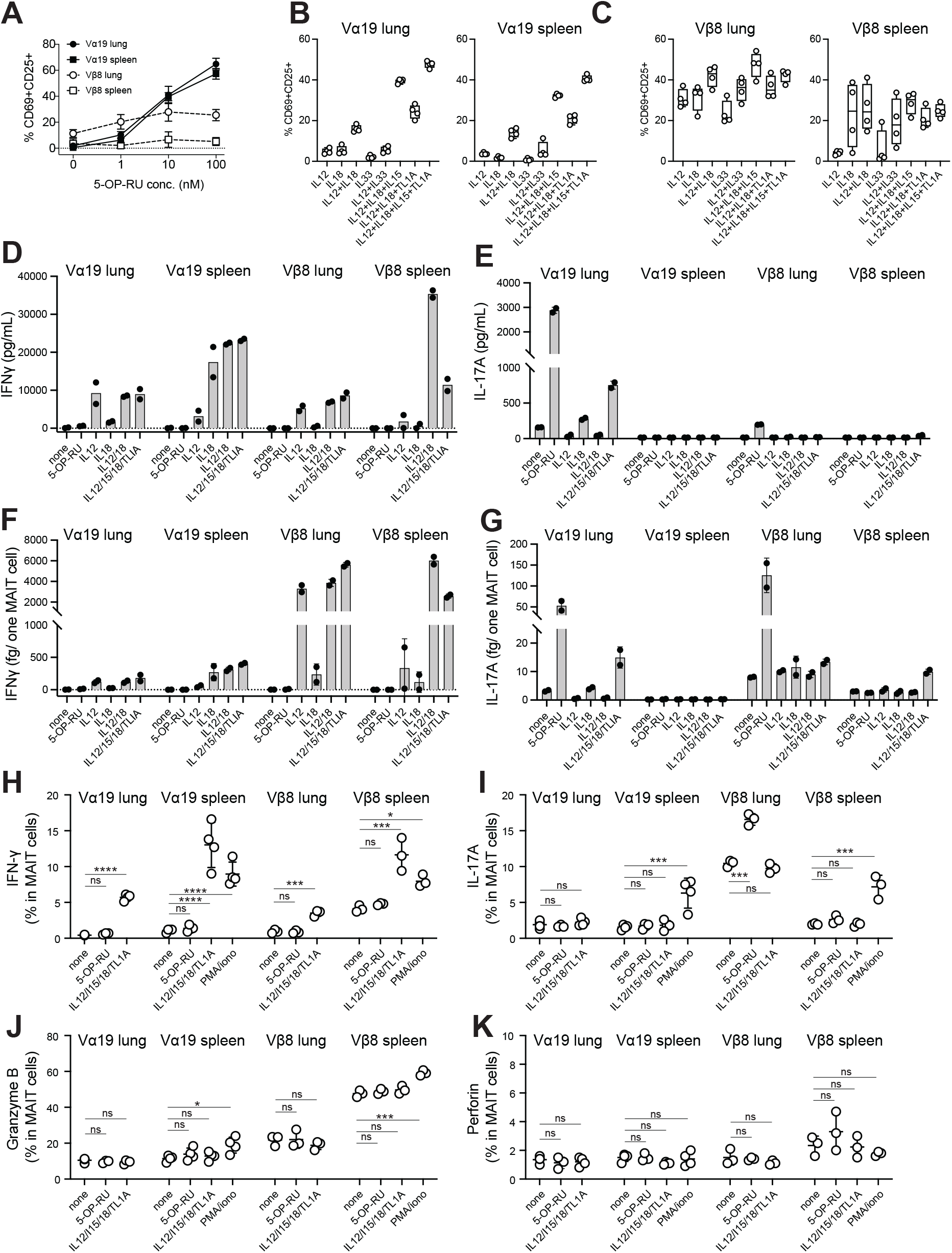
MAIT cell activation and production of IFN-γ and IL17A. (A) Activation of MAIT cells by 5-OP-RU. The percentage of CD69^+^CD25^+^ cells in MAIT cells among lung mononuclear cells and the spleen cells from the indicated mice followed by stimulation with varying amounts of 5-OP-RU (1 - 100 nM) are depicted (n=4 per group). (B and C) Activation of MAIT cells by cytokines. The percentage of CD69^+^CD25^+^ cells in MAIT cells among lung mononuclear cells and the spleen cells from Vα19 mice (Β) and Vβ8 mice (C) followed by stimulation with the combination of the indicated cytokine(s) are depicted (n=4 per group). (D-G) IFN-γ and IL-17A production. Amounts of IFN-γ (D) and IL-17A (E) produced from lung mononuclear cells and from spleen cells upon challenge with the indicated cytokine(s) or 5-OP-RU in Vα19 and Vβ8 mice are shown. IFN-γ (F) and IL-17A (G) produced per cell. The absolute amounts of cytokine produced per cell upon the indicated challenge are indicated. Data from two independent experiments are shown (n=2 per experiment). (H-K) Intracellular cytokine staining. Frequency of MAIT cells expressing IFN-γ (H), IL-17A (I), granzyme B (J) and perforin (K) upon the indicated stimuli is shown (n=4 for Vα19 mouse, n=3 for Vβ8 mouse). * *P*<0.05, *** *P*<0.001, and **** *P*<0.0001 Vα19: Vα19 mouse, Vβ8: Vβ8 mouse

Subsequently, TCR-independent activation of MAIT cells from the lung and spleen of Vα19 mice was assessed with cytokines such as IL-12, IL-18, IL-15, IL-33, and TL1A (Leng et al., 2019). Stimulation with IL-12 or IL-18 alone resulted in little activation, while combination of the cytokines resulted in activation, which was further enhanced by the addition of IL-15 or TL1A (Figure 5B). However, IL-33 failed to induce activation or enhance other cytokine-induced activation in Vα19 mice (Figure 5B). In contrast, Vβ8 mouse lung MAIT cells, exhibiting an activated phenotype without stimulation, were further activated when stimulated with IL-12 or IL-18 alone (Figure 5C). The combination of IL-12/IL18 or IL-12/IL-18/IL-15 led to the highest activation, but the addition of TL1A was not effective in boosting activation as seen in MAIT cells from Vα19 mice (Figure 5C). While activation of Vβ8 mouse spleen MAIT cells fluctuated among individual samples when stimulated with a single cytokine, it was minimized when stimulated with multiple cytokines (Figure 5C) (Ussher et al., 2014, Jesteadt et al., 2018, Leng et al., 2019). We then explored whether activation resulted in the production of cytokines such as IFN-γ and IL-17A. In both Vα19 and Vβ8 mice, stimulated with IL-12 alone or in combination with IL18 and /or TL1A led to IFN-γ production in lung and spleen cells, however, IL-12 or IL-18 alone barely produced FN-γ from Vβ8 mouse spleen cells (Figure 5D). In contrast, 5-OP-RU and the combination of IL12, IL-15, IL-18 and TL1A stimulation engendered significant IL-17A production from the Vα19 mouse lung cells. A similar tendency was observed with Vβ8 mouse lung cells in response to 5-OP-RU (Figure 5E).

Further analysis revealed that the amount of IFN-γ produced per MAIT cell in Vβ8 mice was greater than in Vα19 mice, irrespective of the tissue (Figure 5F). A similar tendency was observed for IL-17A production, except for 5-OP-RU (Figure 5G).

To further ascertain the production of IFN-γ and IL-17A from MAIT cells per se, we examined the potential by intracellular staining. It turned out that ∼10 % of MAIT cells in Vα19 and Vβ8 mouse spleen produced IFN-γ upon PMA/ionomycin or IL-12+IL-18+IL-15+ TL1A, while ∼6% of spleen MAIT cells produced IL-17A upon PMA/ionomycin (Figures 5H-I). Similarly, PMA/ionomycin enhanced the frequency of granzyme B (an effector molecule required for cytotoxic activity)-expressing MAIT cells in Vα19 and Vβ8 mouse spleen, while perforin expression was not enhanced (Figures 5J-K).

The data indicated that MAIT cells in Vα19 and Vβ8 mouse had a potential to produce IFN-γ, IL-17A and granzyme B.

### Tumor resistance in Vβ8 mice

Our previous study demonstrated that MAIT-like cells prepared from iPSCs exert anti-tumor activity, increasing mouse survival upon adoptive transfer (Sugimoto et al., 2022). Therefore, we explored whether the Vα19 and Vβ8 mice could exhibit a similar tumor resistance. Inoculation of Lewis lung carcinoma (LLC) into Vβ8 mice led to an increased mouse survival time relative to control mice, an effect not observed in Vα19 mice (Figure 6A). It was possible that MAIT cells in Vα19 mice were defective with respect to anti-tumor activity. Therefore, we examined whether the adoptive transfer of MAIT cells from Vα19 mice into wild-type mice would improve the survival of the LLC-inoculated mice, which it did. This result demonstrated the preservation of intrinsic anti-tumor activity of Vα19 mice MAIT cells (Figure 6B). As the T cell repertoire of Vα19 mice was reduced at the cost of MAIT cell increase (Figure 4), we then examined whether supplementing T cells from wild-type mice could improve the survival of LLC-inoculated Vα19 mice. Given that T cells, as represented by cytotoxic CD8^+^ T cells, are pivotal for antitumor activity, CD3^+^ T cells or CD8^+^ T cells were adoptively transferred into Vα19 mice. The survival curve upon LLC inoculation for control and Vα19 mice were similar, however, Vα19 mice with the adoptive transfer of CD3^+^ T cells or CD8^+^ T cells from the control mice experienced increased survival time (Figure 6C). These data demonstrate that a simple increase in MAIT cells was not sufficient for survival extension, and T cell supplements, CD8^+^ T cells, in particular, improved the survival rate of Vα19 mice over the control, further highlighting the role of MAIT cells in tumor immunity.

**Figure 6.**
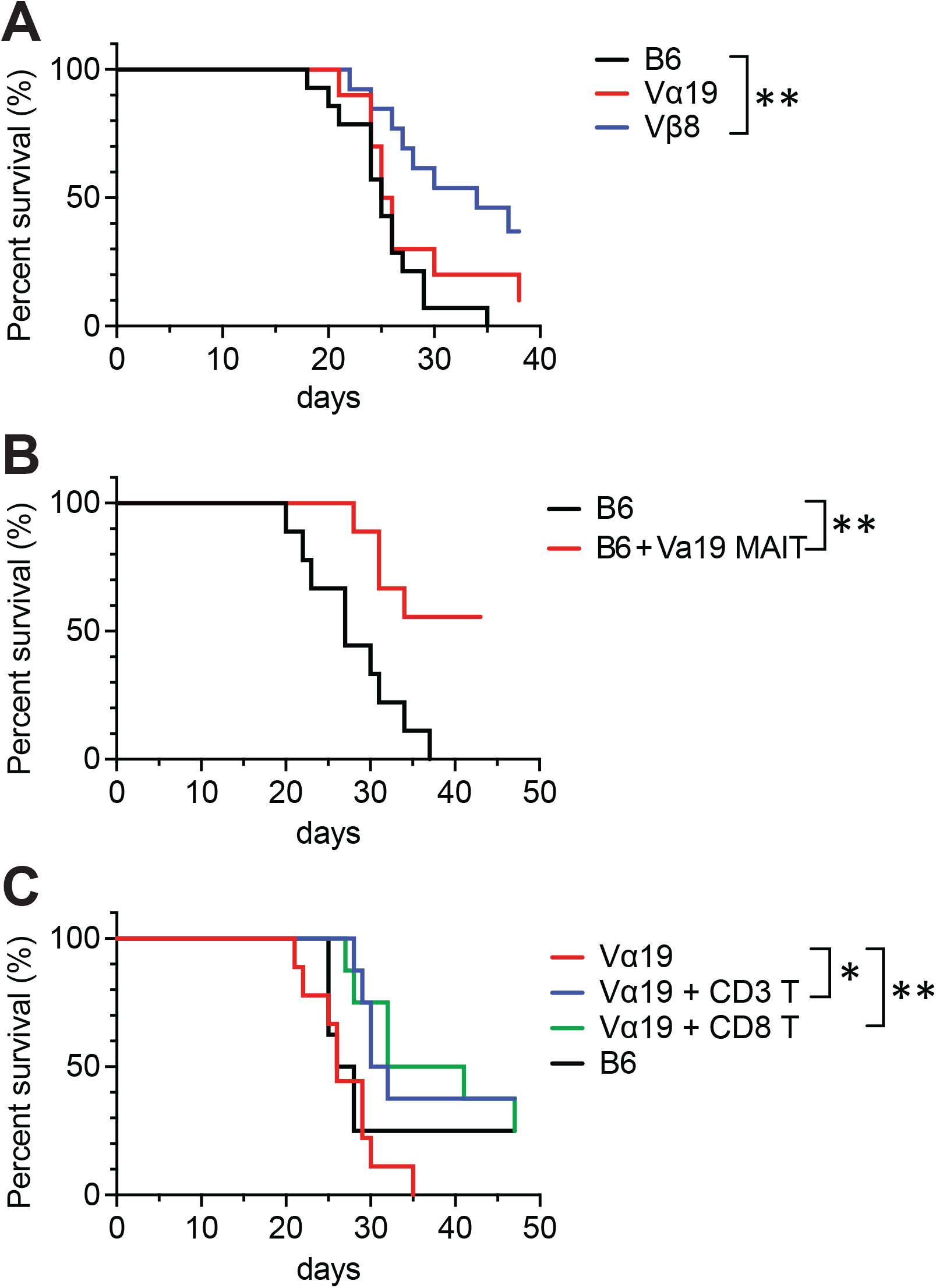
Tumor resistance of Vα19 and Vβ8 mice. (A) Superiority of Vβ8 mice in conferring survival extension. Survival of the indicated strains of the mice was monitored upon tumor inoculation (3 x10^5^ LLC, I. V.) ** *P*<0.01 (n= 14 per strain, log-rank test) (B) Survival superiority conferred by Vα19 MAIT cells. Survival of the control mice (B6) and of the control mice that received MAIT cells prepared from Vα19 mice (B6 +Vα19 MAIT) were monitored as per (A). ** *P*<0.01 (n= 10 per group, log-rank test) (C) Survival superiority conferred to Vα19 mice by CD8^+^ T cells. Survival of the control mice (B6), Vα19 mice (Vα19), and Vα19 mice that received either CD3^+^ T cells (Vα19 + CD3 T, 1.0 x10^6^ cells.) or CD8^+^T cells (Vα19 + CD8 T 1.0 x10^6^ cells) from the control mice was monitored as per (A). * *P*<0.05, ** *P*<0.01 (n= 10 per group, log-rank test)

## Discussion

In this study, we created novel mice models to facilitate the functional analysis of MAIT cells. Previous mouse models for studying the functions of MAIT cells were limited to MR1 knockout (KO) and/or MAIT TCR transgenic mice (Okamoto et al., 2005, Martin et al., 2009, Croxford et al., 2006, Kawachi et al., 2006). While the former allows study on mice devoid of MAIT cells, such mice also lack other MR1-restricted immune cells, making physiological changes difficult to ascribe to MAIT cells alone. Moreover, although data from MR1 KO mice are generally compared to those from wild-type mice, interpretation of the data is often complicated and unconvincing as laboratory mice produce few MAIT cells. Transgenic mice overexpressing TCR specific for MAIT cells can overcome this inherent problem but still suffer from aberrant MAIT cell function, and/or the absence of other T cells (Godfrey et al., 2019). In this respect, it is interesting to note that CAST/EiJ, a strain expressing more MAIT cells than other laboratory strains, exhibited diminished *Mycobacteria tuberculosis* virulence (Dey et al., 2022). Furthermore, Cast/B6 mice have been generated by crossing CAST/EiJ with C57BL/6 to overcome this problem of MAIT cell paucity, although the effect on disease remains unknown (Cui et al., 2015).

In addition to these mice, we herein reported novel mice derived from iPSCs generated from lung MAIT cells. They were characterized by the presence of the rearranged *Tcra* or *Tcrb* locus in the allele, both of which increased the expression of MAIT cells when inherited separately. Although the frequency of MAIT cells in Vα19 mice was higher than in Vβ8 mice, flow cytometric and TCR repertoire analyses revealed the presence of many other T cells including iNKT cells and γδT cells in Vα19 mice (Figure 2 and 4). As *Trav1* is located at the most distal region of the TCRα locus, once MAIT cell-specific TCRα (*Trav1-Traj33*) was selected, iNKT cell-specific TCR (*Trav11-Traj18)* and/or any other *Trav-Traj* rearrangement could not have occurred in the same allele. Thus, it is plausible that V-J recombination from the other allele is at work in these mice to ensure TCRα chain diversity to a certain degree, thus allowing iNKT cell and γδ T cell development.

While CDR3α in thymic MAIT cells from Vα19 mice showed a single sequence, Vβ8 mice exhibited sequence diversity and V-J combination was not confined to *Trav1-Traj33*, counter to previous studies (Figure 4C and Table S2) (Reantragoon et al., 2013, Rahimpour et al., 2015). This unexpected CDR3α diversity may reflect the presence of pre-MAIT cells in the thymus, and the survival or selection of only a fraction before and/or after their egression from the thymus. Furthermore, the abundance of *Trav1-Traj33* transcript, a major combination for MAIT TCRα chain, in mMR1-tet negative T cells described as “non-MAIT T cells” from Vα19 mice might suggest that other MAIT cell subsets, not recognized by 5-OP-RU loaded mMR1-tet, were enriched in the thymus. Alternatively, such an increase might reflect overexpression of rearranged *Trav1-Traj33*, some of which could not be recognized by 5-OP-RU-loaded mMR1-tet due to stoichiometric constraints.

Since the expression of PLZF is essential for proper development and function of MAIT cells, our data indicate that MAIT cells in Vα19 and Vβ8 mice were functional (Figure 1E). Furthermore, expression of RORγt and/or T-bet in MAIT cells concomitant with the production of IL-17A and/or IFN-γ further underpins the notion that MAIT cells in these mice were authentic, and distinguished from those found in Vα19-Cα^-/-^ transgenic mice (Figures 1E, Figures 5D-I, and S2) (Rahimpour et al., 2015).

While the distribution of MAIT1 and MAIT17 in Vα19 mice and Vβ8 mice were similar to that in C57BL/6, it has yet to be analyzed MAIT cells across the tissues by a single cell RNA-sequencing. This could further underpin the above notion and shed much light on the tissue-specific MAIT cell subset distribution and function.

Regardless of the increase in MAIT cells concomitant with T cells harboring V-J combinations other than *Trav1-Traj33* in Vα19 mice, it was not sufficient to fully recover the frequency of CD8^+^T cells. In particular, a decrease of CD8^+^T cells in Vα19 mice might directly lead to compromised anti-metastatic activity (Figure 5A). While our previous study demonstrated that NK cells are an important partner for MAIT cells to exert anti-metastatic function, this study further implied that CD8^+^T cells are required (Sugimoto et al., 2022). In this respect, the influence of CD8^+^T cells on MAIT cells in tumor immunology warrants further study. Given the frequency of CD4^+^ T cells was barely affected in Vα19 mice, future studies should examine the mechanism(s) by which *Trav1-Traj33* in the allele engendered a decrease in CD8^+^ T cells. As MR1 belongs to MHC Ib, it is tempting to postulate that there is competition between MHC I and MHC Ib, but not MHC II and MHC Ib in T cell development.

### Limitations of the study

Though we have reported mice with increased MAIT cell numbers, equivalent to or superior to that of humans, without genetic modification and their utility in cancer immunology, not all MAIT cell functions in Vα19 and Vβ8 mice have been investigated. In particular, the effects of allelic exclusion in the presence of the rearranged *Trbv* on MAIT cell activation, cytokine production, and cytolytic activity in Vβ8 mice are yet to be examined. Similarly, the nature of the immune cells including MAIT cells in Vα19 mice should be further explored.

MAIT cells from Vα19 mice exhibited anti-metastatic properties upon adoptive transfer, however, it remains undetermined whether MAIT cells were composed of a unique population, or of different cell subsets with diverse functions and different developmental stages. Moreover, tissue-specific MAIT cell characterization is fundamental for clarifying MAIT cell function in each tissue and in understanding the integral role of MAIT cells in health and disease.

## Author contribution

H. W., C. S., and H. F. designed experiments and provided conceptual input. C.S and H.W. performed experiments, analyzed data, and wrote the manuscript.

## Acknowledgments

We thank Y. Murakami and M. Ohyama (Dokkyo Medical University) for technical help, Dr. H. Kon, Y. Machida, and T. Tsukahara (Animal facility, Dokkyo Medical University) for tumor experiments, Drs. S. Nishioka and M. Ikawa (Biken, Osaka University, Osaka, Japan) for chimeric mice, the NIH Tetramer core facility (Emory University, GA, USA) for mMR1-tetramers, and Advanced Animal Model Support for supporting chimeric mice (JSPS KAKENHI 16H06276). This work was supported by The Science Research Promotion Fund 2018-2019 (The Promotion and Mutual Aid Corporation for Private Schools of Japan) and 21K19732 (JSPS KAKENHI) to H.W., 17H03565 (JSPS KAKENHI) and 26430084 (JSPS KAKENHI) to C. S., and 20K10435 (JSPS KAKENHI) to H.F.

## RESOURCE AVAILABILITY

### Lead Contact

Further information and requests for resources and reagents should be directed to and will be fulfilled by the Lead Contact, Hiroshi Wakao

### Materials Availability

The mouse strains (Vα19 and Vβ8) generated in this study will be made available to qualified investigators by having Material Transfer Agreement.

### Data and Code Availability

This study did not generate any unique code or database. All data supporting the findings of this study are available within the paper and its supplemental information.

## EXPERIMENTAL MODEL AND SUBJECT DETAILS

### Mice

All mouse experiments were performed in accordance with approval from the Institutional Animal Care and Use Committee of Dokkyo Medical University. Mice were housed in the Animal Research Center, Dokkyo Medical University, under specific pathogen free conditions with controlled lighting and temperature with food and water provided *ad libitum*. For breeding, male and female mice aged between 8 to 30 weeks were used. For tissue cell analyses and cancer models, male and female mice aged between 8 to 12 weeks were used.

### Cell lines

Mouse cancer cell line LLC were cultured in DMEM supplemented with 10% FBS at 37ºC in 5% CO_2_.

## METHOD DETAILS

### Generation of chimeric mice from MAIT-iPSCs

iPSCs were established previously from lung MAIT cells of male C57BL/6NJcl (Sugimoto et al., 2022). MAIT-iPSC clones L7-1, L11-1, and L19-1 were selected by karyotype analysis (80-90% euploid metaphases) and microinjected into ICR 8-cell stage embryos at NPO for Biotechnology Research and Development, Osaka University (Osaka, Japan).

### Generation of Vα19 and Vβ8 mice from the chimeric mouse

To confirm germline transmission of an allele derived from MAIT-iPSCs, the male chimeras were crossed with C57BL/6NJcl females (CLEA Japan) and progenies were genotyped for the rearranged TRA (*Trav1-Traj33*) and TRB (*Trbv13-3-d1-j1-2* or *Trbv19-1-d2-j2-3)* loci for MAIT cells by PCR. Offspring of a germline-transmitted chimera (#26) harboring *Trav1-Traj33* or *Trbv13-3-d1-j1-2* in an allele were crossed to obtain homozygous mice for the rearranged locus, and the obtained mice were further crossed with C57BL/6Jcl (CLEA Japan) to generate hemizygous mice harboring either *Trav1-Traj33* (Vα19 mice) or *Trbv13-3-d1-j1-2* (Vβ8 mice). Vα19/Vβ8 mice were hemizygous for both *Trav1-Traj33* and *Trbv13-3-d1-j1-2*.

### Mouse genotyping

Ear punches from mice were treated in 50mM NaOH at 95ºC for 10 min and centrifuged at 15000 rpm for 10 min. The supernatant was used as genomic DNA sample after neutralization. Primers were designed to detect the rearranged configuration of TRA and TRB specific for MAIT-iPSC using Primer-BLAST (Figure S1). PCR amplification was performed with the specific primers and OneTaq master mix (New England Biolabs) under the following condition: 94ºC for 30 sec, 35 cycles of 94ºC for 30 sec, 55ºC for 30 sec, and 68ºC for 30sec, and 68ºC for 3 min. PCR products were analysed on 2% agarose gel electrophoresis.

### Cell isolation from tissues

#### Spleen, thymus, and lymph nodes

Cell suspension was prepared by mashing tissues through a 40-µm mesh cell strainer with a syringe plunger. Single cells were suspended in RPMI1640 supplemented with 10% FBS, 10 mM HEPES pH 7.0, 0.1 mM 2-mercaptoethanol, and 100 IU/ml of penicillin/100 µg/ml streptomycin (referred cR10) and spun down at 400×*g* for 4 min. The cell pellet was suspended in sterile ice-cold MilliQ water for 15 sec to lyse red blood cells, and immediately neutralized with an equal volume of 2× PBS containing 4% FBS. After centrifugation, cells were resuspended in cR10.

#### Lungs and liver

Single-cell suspensions from the lungs and liver were prepared using enzymatic digestion. Briefly, tissues were placed into a GentleMACS C-tube (Miltenyi Biotec) and cut into approximately 5-mm^3^ pieces. Four milliliters of tissue digestion solution (90 U/ml collagenase Yakult, 275 U/ml collagenase type II, 145 PU/ml Dispase II, and 4% BSA in HBSS) was added per tissue and tissues were further homogenized with the GentleMACS dissociator (Miltenyi Biotec) using the following program: m_lung_01_02 for the lungs and m_liver_03_01 for the liver. Suspensions were then incubated at 37ºC for 30 min under gentle rotation, followed by dissociation with m_lung_02_01 for the lungs and m_liver_04_01 for the liver, and subjected to discontinuous density centrifugation over layers of 40% and 60% Percoll at 400×*g* for 20 min. Cells were recovered from the 40%-60% Percoll interface, washed with PBS, and then suspended in cR10.

#### Intestines

The intestines were longitudinally incised, and their contents were thoroughly washed three times with PBS by vigorous shaking. Tissue dissected into 1-cm pieces were placed into a 50-ml conical tube and washed vigorously by shaking three times with PBS. After discarding the supernatant, tissues were treated with 40 ml of intraepithelial lymphocyte (IEL)-washing solution (HBSS containing 1 mM DTT, 5 mM EDTA, and 1% BSA) by shaking vigorously at 37ºC for 30 min. After washing three times with MACS buffer (PBS containing 2 mM EDTA and 0.5 % BSA), tissue was further washed with 40 ml HBSS. After removal of the supernatant, tissues were placed into a GentleMACS C-tube (Miltenyi Biotec), cut into small pieces with scissors, and 4 ml of the Tissue digestion solution (see above) was added. Tissues were processed using the following program: m_brain_01_02 and digested at 37ºC for 30 min under gentle rotation followed by dissociation with the program: m_intestine_01_01. Cell suspensions were subjected to Percoll discontinuous density centrifugation and isolated cells were resuspended in cR10 for further analysis.

### Flow cytometry

Cells were stained with the antibodies listed in STAR methods. For analyses of cell surface marker staining, 7-AAD was used to discriminate between live and dead cells. For expression analysis of the transcription factors in MAIT cells, the cells were stained with anti-TCRβ and mMR1-tet, then fixed and permeabilized with Transcription Factor buffer set (BD Biosciences) according to the manufacture’s instruction. Then, the cells were stained with anti-PLZF, -T-bet, and -RORγt antibodies. For intracellular cytokine staining, the cells were treated with a different combination of the cytokines, Cell Stimulation Cocktai (ThermoFisher), or 5-OP-RU for 15 h. Protein Transport Inhibiitor Cocktail (ThermoFisher) was added 1.5 h after the start of stimulation. Thereafter, the cells were subjecte to intracellular staining with BD Cytofix/Cytoperm buffer kit according to the manufactures’s instruction (BD BioScience). Cells were analyzed with the MACSQuant cell analyzer (3 lasers, 10 parameters, Miltenyi Biotec) or the AttuneNxT acoustic focusing cytometer (4 lasers, 14 parameters, Thermo-Fisher Scientific). Data were processed using FlowJo software (ver.9 or 10, BD Biosciences). Cell sorting was performed using a FACSJazz cell sorter (2 lasers, 8 parameters, BD Biosciences).

### MAIT cell activation

Cells from lungs (5x10^5^ cells/well) or spleens (1x10^6^ cells/well) of Vα19 and Vβ8 mice were stimulated with cytokines (200 ng/ml each) or 5-OP-RU (1-100 nM) in 96-well culture plates at 37ºC, 5% CO_2_ for 18 h. The culture supernatants were collected for cytokine assay described below and the cells were stained to determine MAIT cell activation by flow cytometry.

### Cytokine production

The quantification of cytokines was performed with the LegendPlex mouse Th cytokine panel according to the protocol provided by the manufacturer (Biolegend, USA). The amounts of cytokine per cell were calculated as follows: total amounts of the cytokine/the number of MAIT cells (mMR1-tet^+^TCRβ^+^ cells) in the indicated source.

### TCR repertoire analysis

MAIT cells (mMR1-tet^+^TCRβ^+^ cells) and non-MAIT T cells (mMR1-tet^-^CD4^+^CD8^+^TCRβ^+^ cells) were sort-purified from thymocytes of Vα19 and Vβ8 mice and stored into RNAlater (Qiagen). Total RNA was isolated using RNeasy Mini kit (Qiagen) and resultant RNAs (10 ng per sample) were subjected to cDNA library construction (SMARTer Mouse TCR a/b profiling kit, Takara Bio, Japan). The next generation sequencing was performed with MiSeq (Illumina, USA) at FASMAC (Kanagawa, Japan). Sequencing data were aligned to reference mouse TCR V/D/J sequences registered in ImMunoGeneTics database and then assembled into TCR clones using MiXCR-3.0.5 (Bolotin et al., 2015) and TCR repertoire data were visualized using VDJtools-1.2.1 (Shugay et al., 2015) at ImmunoGeneTeqs, Inc (Chiba, Japan).

### Tumor resistance

Lewis lung carcinoma (LLC) suspended in HBSS was intravenously (I.V.) inoculated (3.0 × 10^5^ cells/mouse) into the indicated mouse strains. For MAIT cell supplementation experiments, MAIT cells were isolated from lungs and spleens of Vα19 mice. In brief, cells were stained with APC-labeled mMR1-tet and mMR1-tet positive cells were isolated with anti-APC microbeads and LS columns (Miltenyi Biotec). The isolated MAIT cells were intraperitoneally (IP) injected (1.0 × 10^6^ cells/mouse) into control (C57BL/6) mice 3 days before LLC inoculation. For CD3^+^T cells and CD8^+^T cells supplement experiments, CD3^+^ T cells were purified from C57BL/6 spleen cells with MojoSort mouse CD3 T cell isolation kit (Biolegend, USA), while CD8^+^T cells were isolated by adding biotin-labeled anti-CD4 antibody into MojoSort mouse CD3 T cell isolation kit. The cells purified were IP injected (3.0 × 10^5^ cells/mouse) into Vα19 mice 3 days before LLC inoculation. In the survival assay, mice were considered to be dead when they showed a humane end-point, such as acute weight loss, hypothermia, and severe gait and/or consciousness disturbance, and the survival time of mice was plotted as a Kaplan-Meier curve.

## QUANTIFICATION AND STATISTICAL ANALYSIS

Statistical analyses were conducted using Prism 9.9.6 for macOS (GraphPad). In each experiment, deta variability was shown as standerd diviation (SD). One-way ANOVA was used for analysis of the intracellular cytokines and the effector molecules. The Log-rank test was used for survival analyses between the two indicated groups.

### Legends for supplemental tables

**Table S1 (related to Figure 4) TCR**α **repertoire for thymic MAIT cells from V**α**19 mice**

The absolute count of clones carrying the indicated V and J is shown with the CDR3α sequences in nucleotide (nt) and amino acid (aa) concomitant with the relative frequency (freq).

**Table S2 (related to Figure 4) TCR**α **repertoire for thymic non-MAIT T cells from V**α**19 mice**

The absolute count of clones carrying the indicated V and J is shown with the CDR3α sequences in nucleotide (nt) and amino acid (aa) concomitant with the relative frequency (freq).

**Table S3 (related to Figure 4) TCR**β **repertoire for thymic MAIT cells from V**α**19 mice**

The absolute count of clones carrying the indicated V and J is shown with the CDR3β sequences in nucleotide (nt) and amino acid (aa) concomitant with the relative frequency (freq).

**Table S4 (related to Figure 4) TCR**β **repertoire for thymic non-MAIT T cells from V**α**19 mice**

The absolute count of clones carrying the indicated V and J is shown with the CDR3β sequences in nucleotide (nt) and amino acid (aa) concomitant with the relative frequency (freq).

**Table S5 (related to Figure 4) TCR**α **repertoire for thymic MAIT cells from V**β**8 mice**

The absolute count of clones carrying the indicated V and J is shown with the CDR3α sequences in nucleotide (nt) and amino acid (aa) concomitant with the relative frequency (freq).

**Table S6 (related to Figure 4) TCR**α **repertoire for thymic non-MAIT T cells from V**β**8 mice**

The absolute count of clones carrying the indicated V and J is shown with the CDR3α sequences in nucleotide (nt) and amino acid (aa) concomitant with the relative frequency (freq).

**Table S7 (related to Figure 4) TCR**β **repertoire for thymic MAIT cells from V**β**8 mice**

The absolute count of clones carrying the indicated V and J is shown with the CDR3β sequences in nucleotide (nt) and amino acid (aa) concomitant with the relative frequency (freq).

**Table S8 (related to Figure 4) TCR**β **repertoire for thymic non-MAIT T cells from V**β**8 mice**

The absolute count of clones carrying the indicated V and J is shown with the CDR3β sequences in nucleotide (nt) and amino acid (aa) concomitant with the relative frequency (freq).

## KEY RESOURCES TABLE

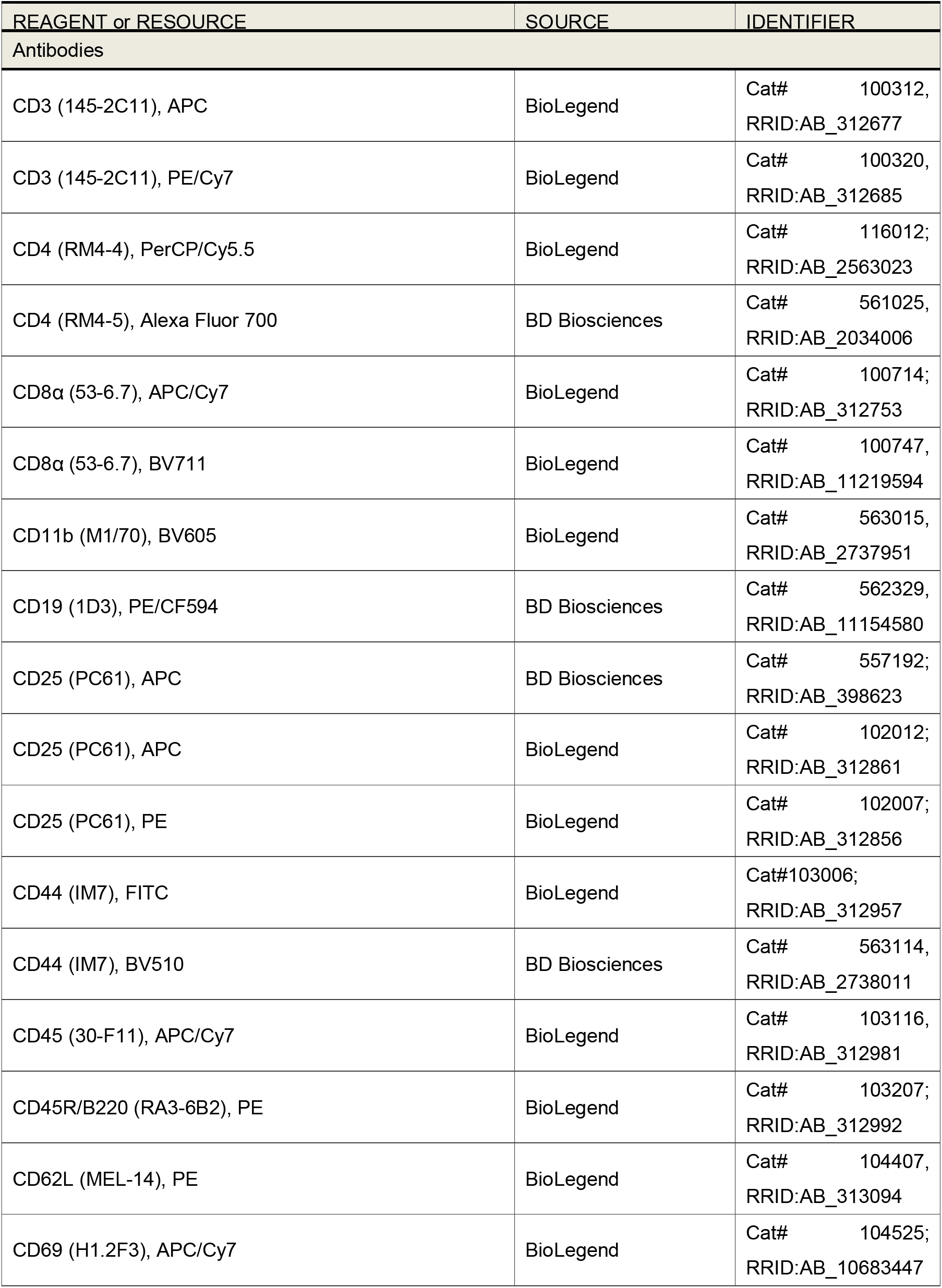
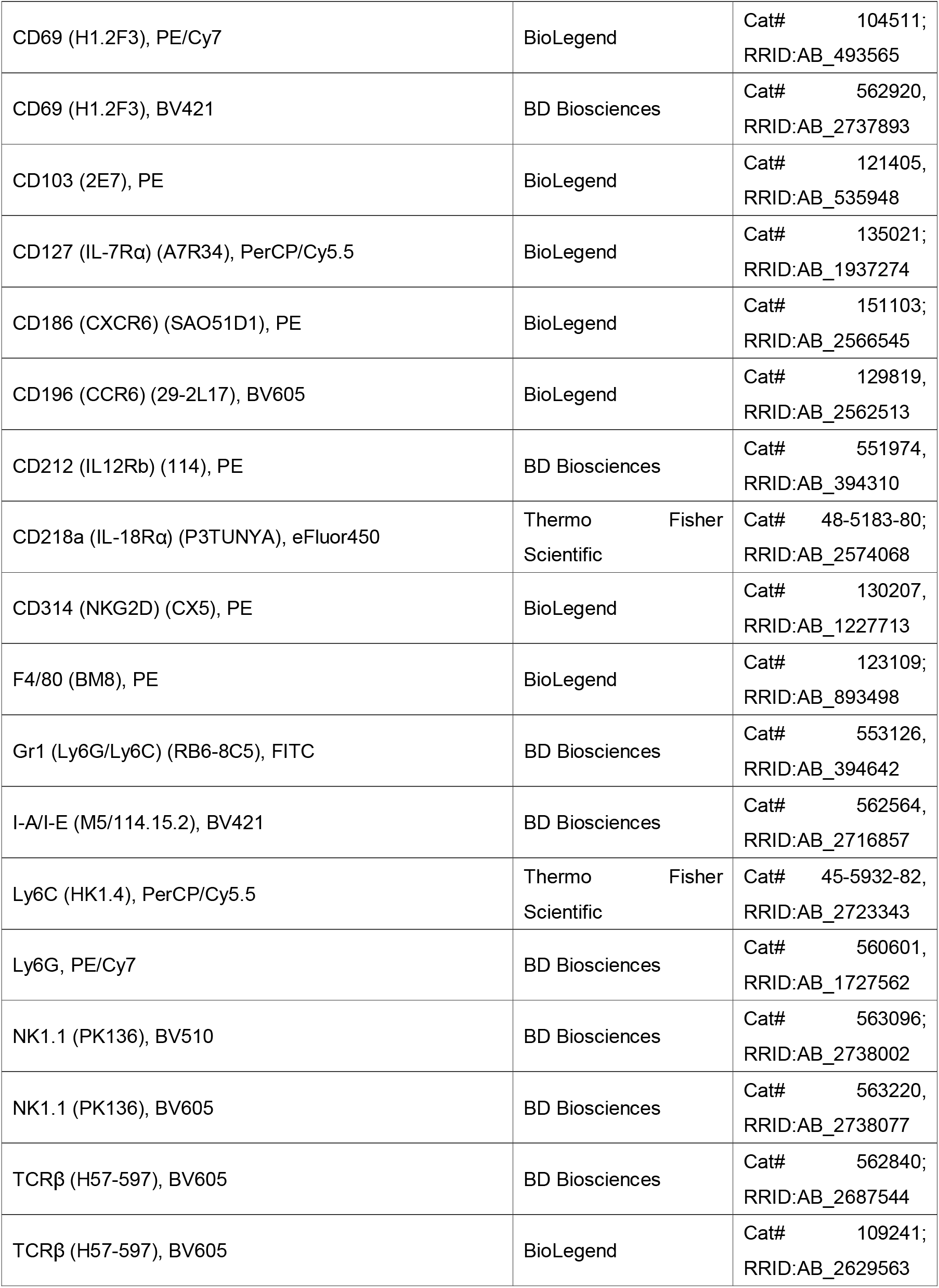
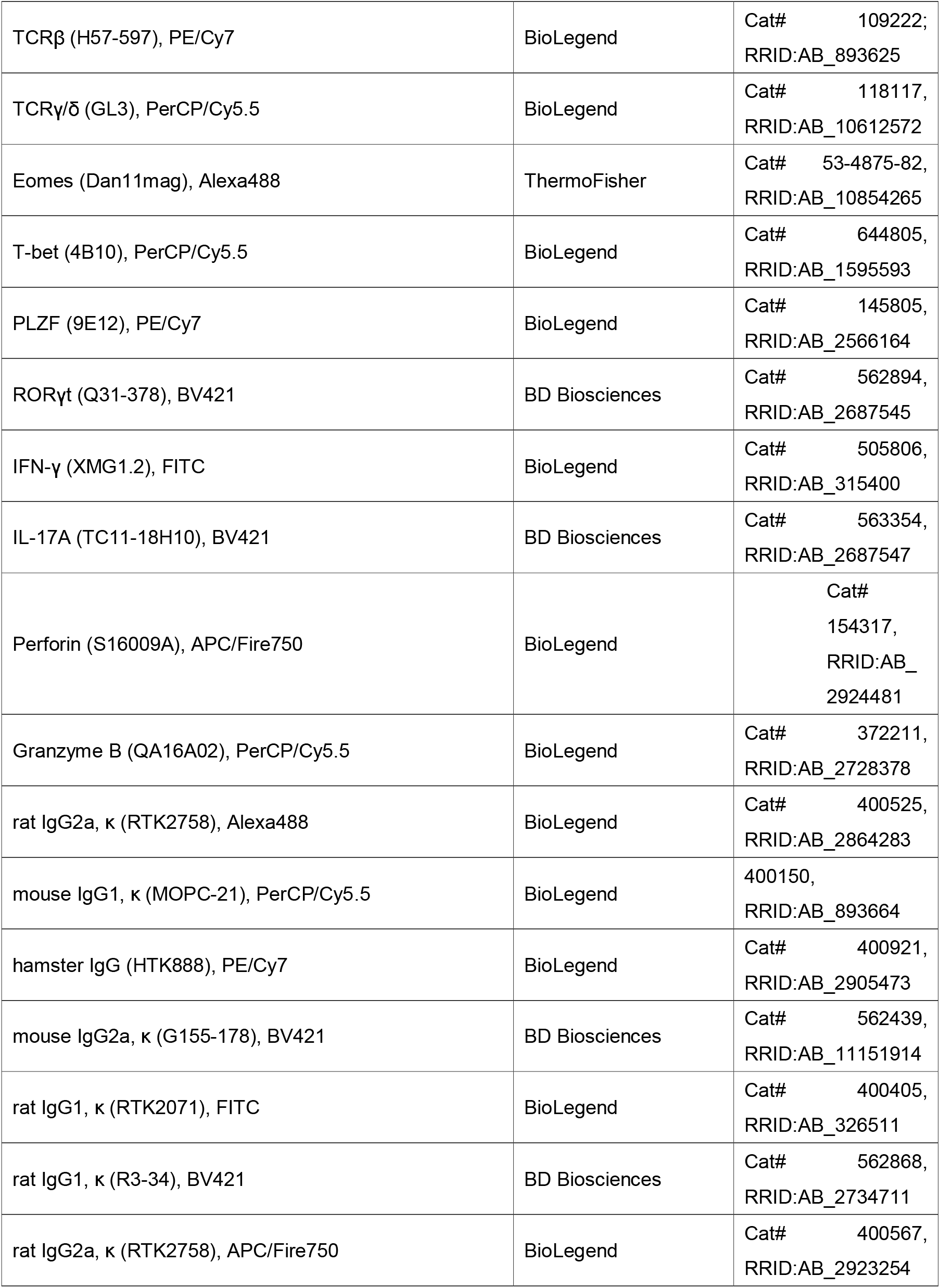
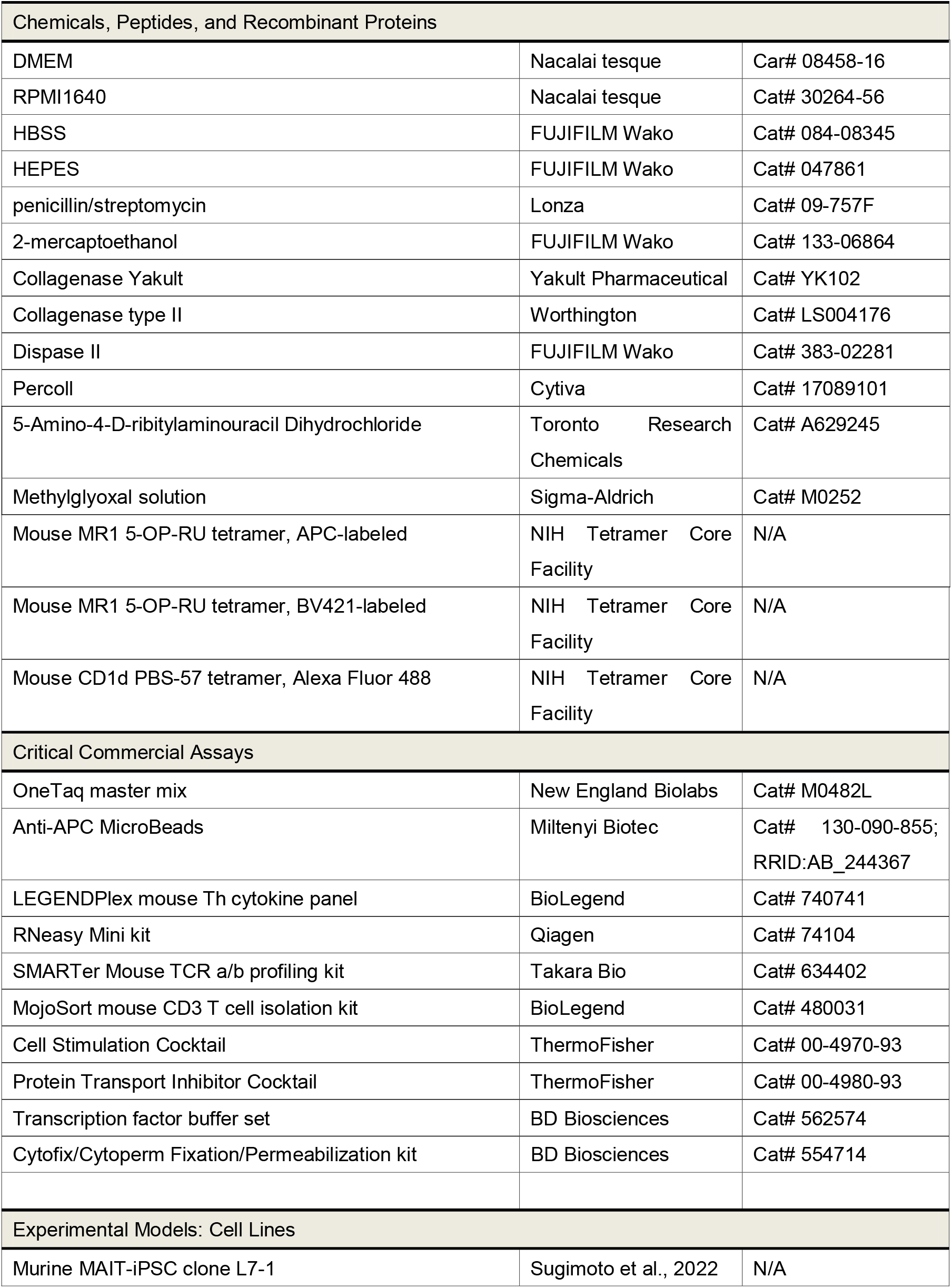
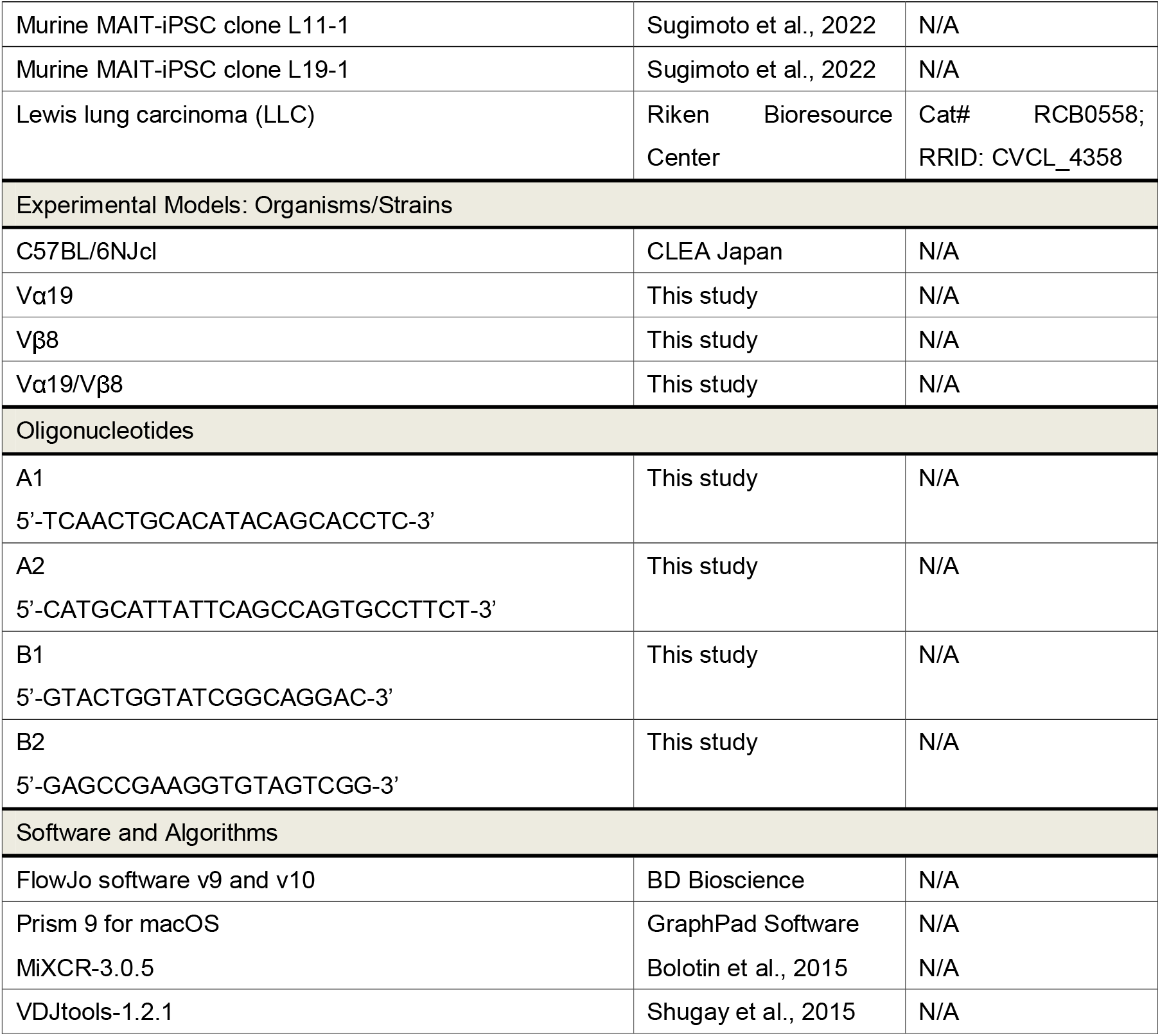

