## Supplementary material for "Mice generated with induced pluripotent stem cells derived from mucosal-associated invariant T cells": Fig. S1

### Figure S1

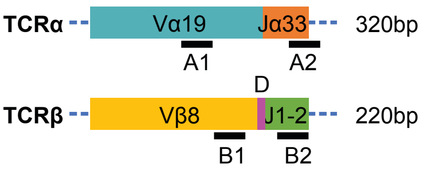


A1: 5’-TCAACTGCACATACAGCACCTC-3’
A2: 5’-CATGCATTATTCAGCCAGTGCCTTCT-3’
B1: 5’-GTACTGGTATCGGCAGGAC-3’
B2: 5’-GAGCCGAAGGTGTAGTCGG-3’

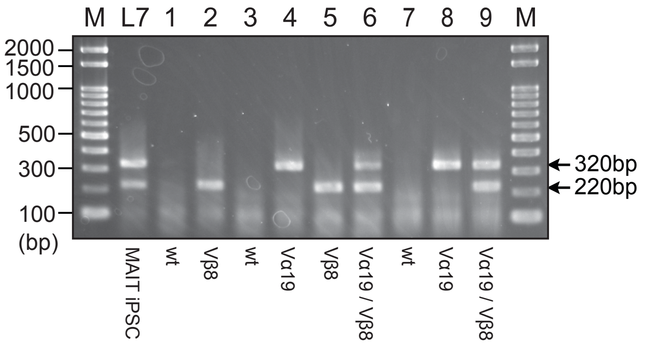


F1 progenies produced from crosses between #26 chimeric mouse and B6 wt mouse.

| | No. of F1 |
|---|---|
| Vα19 | 6 |
| Vβ8 | 14 |
| Vα19/Vβ8 | 4 |
| wt | 12 |
| total | 36 |

**Figure S1 (related to Figure 1) TCR locus configuration in Vα19 mice and Vβ8 mice**
Schematic representation of the rearranged *Trav1* (Vα19) -*Traj33* (Jα33) and *Trabv13-3* (TCRβ8-3) -*Trvd-Trvj* in the alle of Vα19 mice and Vβ8 mice, respectively (upper left panel). Note that Vα19/ Vβ8 mice carry both *Trav1-Traj33* and *Trabv13-3-Trvd-Trvj* in the allele. The position and sequences of the primer sets detecting these configurations are indicated (upper panels). Representative PCR results to distinguish each strain are shown (MAIT cell-derived iPSC served as positive control for both *Trav1-Traj33* and *Trabv 13-3-Trvd-Trvj*). Summary of the crossing between the chimeric mouse generated from MAIT cell-derived iPSC (#26) and C57BL/6 females. The number of F1 carrying the indicated genotype is depicted.
