## Supplementary material for "Mice generated with induced pluripotent stem cells derived from mucosal-associated invariant T cells": Fig. S2

### Figure S2

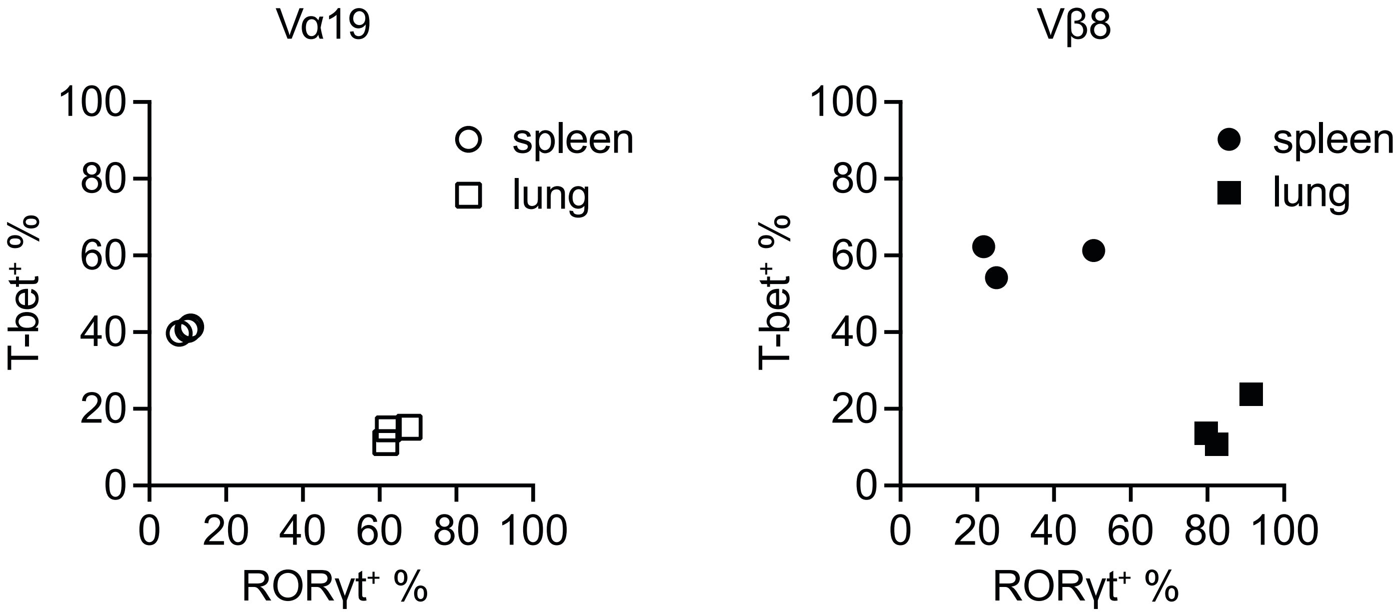


**Figure S2 (related to Figure 1) Correlation between MAIT1 and MAIT17 in spleen and lung**
Scatter plot showing the percentage of lung and spleen MAIT cells expressing T-bet (MAIT1) and/or RORγt (MAIT17) in Vα19 mouse (left panel, Vα19) and in Vβ8 mouse (right panel, Vβ 8). (n=3)
