## Supplementary material for "Mice generated with induced pluripotent stem cells derived from mucosal-associated invariant T cells": Fig. S3

### Figure S3

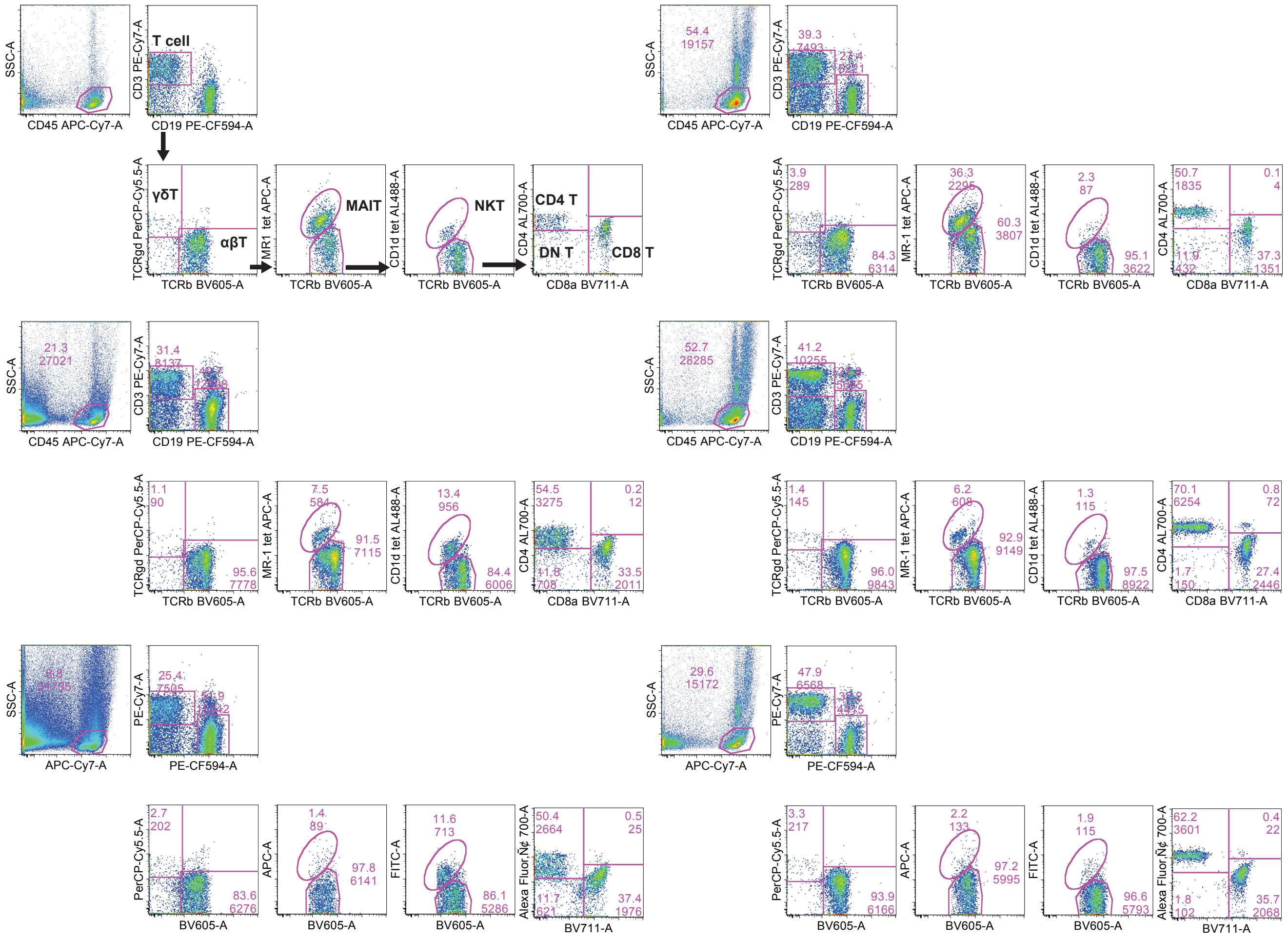


**Figure S3 (related to Figure 2) Gating strategy to identify γδ, MAIT, iNKT, and other T cells**

Blood leukocytes ($CD45^+$ cells) were further gated with the indicated antibodies. Vα19 mice (upper panels), Vβ8 mice (middle panels), and the control mice (lower panels).
