## Supplementary material for "Mice generated with induced pluripotent stem cells derived from mucosal-associated invariant T cells": Fig. S4

### Figure S4

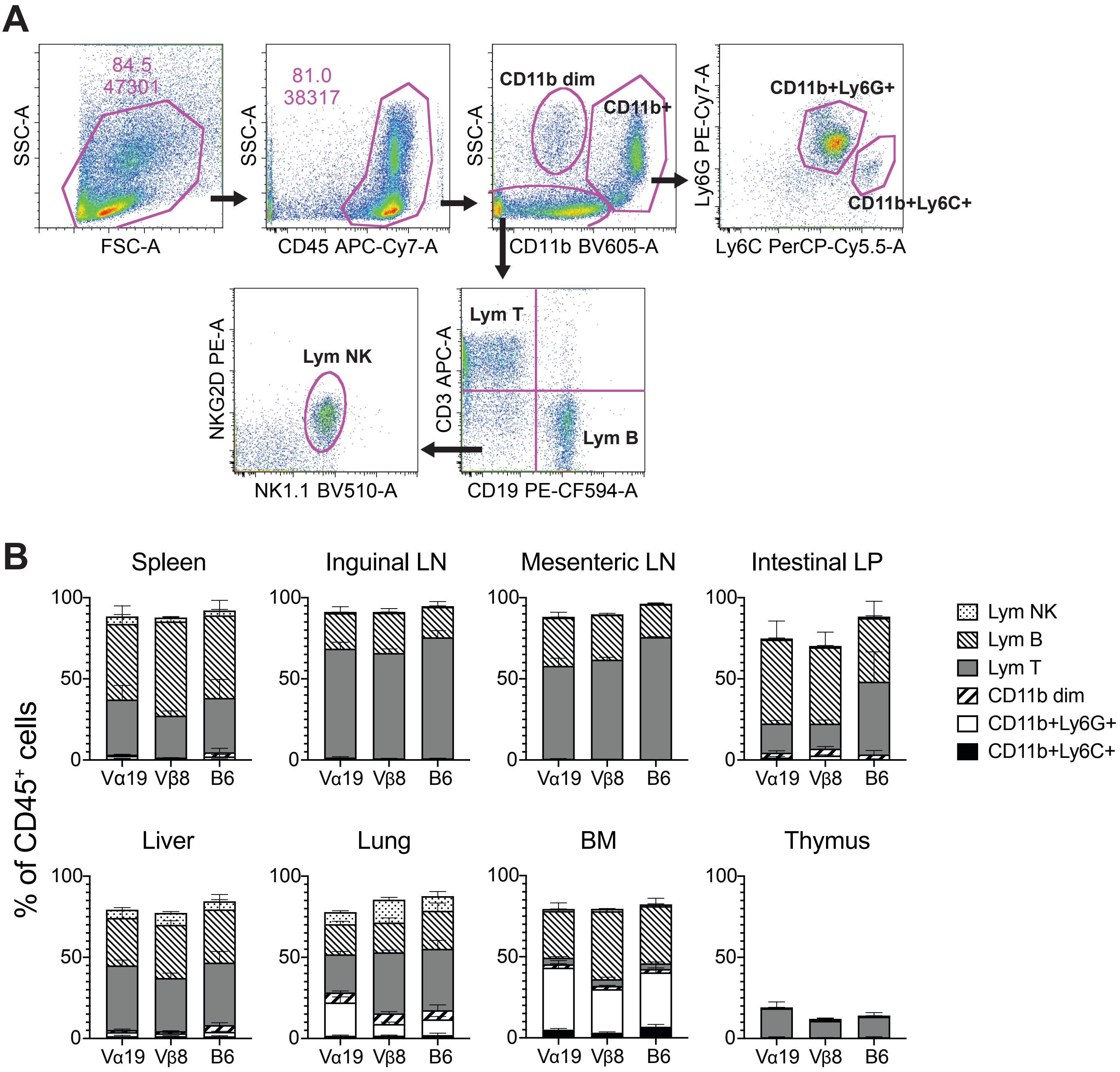


**Figure S4 (related to Figure 2) Various leukocyte populations in tissues from Vα19 and Vβ8 mice**

(A) Gating strategy to identify macrophage, neutrophil, monocytes, NK cells, and T cells. Isolated cells from mouse tissues were analyzed by flow cytometry with the indicated antibodies. CD45$^+$CD11b$^{dim}$ cells were identified as macrophages. CD11b$^+$ cells were classified into CD11b$^+$Ly6G$^+$ (neutrophils) and CD11b$^+$Ly6G$^+$ (monocytes). CD11b negative population was further divided according to the expression of CD3 and CD19. T cells (Lym T) were defined as CD3$^+$CD19$^-$, while B cells (Lym B) were CD3-CD19+. CD3-CD19- population was further explored by expression of NKG2D and NK1.1 and the NKG2D$^+$NK1.1$^+$ cells were identified as NK cells (Lym NK). (B) Relative frequency of NK, B, and T cells, macrophage, neutrophil, and monocytes in tissues from mice. Relative frequency of each subset in the organs is indicated in the graph bar. Data are representative of three independent experiments with a similar tendency. Vα19 mice (Vα19), Vβ8 mice (Vβ8), and control mice (B6).
