## Supplementary material for "Mice generated with induced pluripotent stem cells derived from mucosal-associated invariant T cells": Fig. S5

### Figure S5

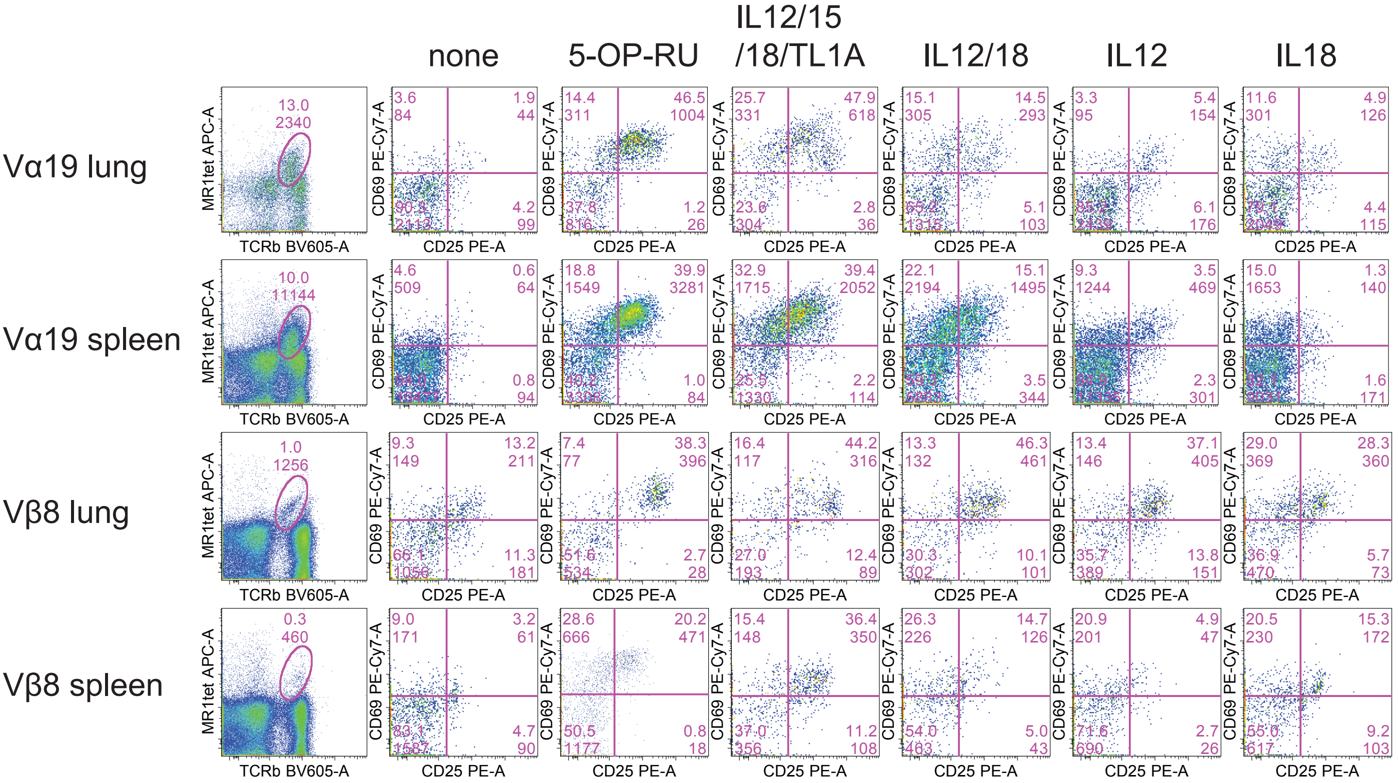


**Figure S5 (related to Figure 5) MAIT cells activation induced by an agonist and cytokine(s)**
Flow cytometric profiles of CD25 and CD69 expression in MAIT cells (TCRβ$^{+}$MR1-tet$^{+}$ cells, the right panels) are shown. Stimuli are indicated above the panels. Data are representative from three independent experiments with a similar profile.
